# MYC/MAX balance dictates cell progenitor fate by altering the HOX program in the *Drosophila* eye

**DOI:** 10.64898/2025.12.12.693907

**Authors:** Sara Monticelli, Giorgio Milazzo, Suleman Khan Zadran, Martina Santulli, Nicola Balboni, Silvia Strocchi, Ettore De Giorgio, Pieter Mestdagh, Angela Giangrande, Roberto Bernardoni, Giovanni Perini

**Affiliations:** Department of Pharmacy and Biotechnology Alma Mater Studiorum - University of Bologna, via Francesco Selmi 3, Bologna, 40126, Italy; University of Ghent, Campus UZ, C. Heymanslaan 10, Blok B (entrance 36), 9000 Ghent, Belgium; IGBMC, Institut de Génétique et de Biologie Moléculaire et Cellulaire, 67400 Illkirch, France; Centre National de la Recherche Scientifique, UMR 7104, 67400 Illkirch, France; Institut National de la Santé et de la Recherche Médicale, UMR, S 1258, 67400 Illkirch, France; Université de Strasbourg, IGBMC UMR 7104- UMR-S 1258, 67400 Illkirch, France; Istituto di Ricerca e Cura a Carattere Scientifico (IRCCS) AOUBO Sant Orsola - Laboratory of Preclinical and Translational Research in Oncology (PRO), Bologna, 40138, Italy; IGBMC, Université de Strasbourg, CNRS UMR 7104, INSERM UMR S 1258, 67400 Illkirch, France; Institut Curie, PSL Research University, CNRS UMR3215, INSERM U934, UPMC Paris-Sorbonne, 75005 Paris, France

**Keywords:** MYC, MYC/MAX ratio, cell transdetermination, HOX genes, *Drosophila*

## Abstract

The transcription factor MYC acts canonically as heterodimer with MAX to promote growth, proliferation, and metabolic adaptation. Although MYC and MAX relative levels are normally kept in balance under physiological conditions, those levels are often dysregulated in many human cancers where MYC is overexpressed and MAX can sometimes be reduced. How altered stoichiometry of MYC/MAX can impact on cell biology is not completely understood. Using the *Drosophila* eye amenable to genetic modifications, we demonstrate that the relative abundance of the fly MYC ortholog dMyc and its partner dMax affect progenitor cell fate. Elevating the dMyc/dMax ratio—through dMyc or human MYCN overexpression, dMax depletion, or combined manipulations—disrupts photoreceptor differentiation and ectopically induces the thoracic HOX gene *Antennapedia* within the eye disc. When such a ratio is driven to an extreme degree, the change in cell fate results in eye-to-wing transdetermination. Conversely, restoring dMax levels rescues these phenotypes, demonstrating that stoichiometry, not absolute MYC levels, shapes the response. Transcriptomic profiling reveals global repression of eye-specific regulators (*e.g., eyegone* and *prospero*) and ectopic activation of wing determinants (*e.g., vestigial* and *nubbin*) while genetic interactions identify the cephalic HOX gene *Deformed* as a key mediator modulating MYC-dependent identity changes. Our findings uncover MYC/MAX imbalance as a developmental switch that can rewire HOX expression and reprogram progenitor identity, suggesting a conserved mechanism underlying MYC-driven cellular plasticity in cancer.

## Introduction

The MYC family of transcription factors (MYC, MYCN, and MYCL) modulates essential cellular processes, including growth, proliferation, metabolism, apoptosis and differentiation, by controlling a broad array of target genes. Dysregulation of MYC proteins, and consequently of these processes, contributes to the initiation and progression of numerous human cancers (Dhanasekaran et al., 2022). Intriguingly, recent evidence highlights a role for MYC proteins in tumor cell transdifferentiation, the direct conversion of one differentiated cell type into another, which contributes to tumor heterogeneity and therapy resistance (Patel et al., 2021), (Quintanal-Villalonga et al., 2021). Although the precise mechanisms underlying MYC-mediated control of cell fate remain elusive, MYC-driven chromatin reshaping can disrupt the normal expression of HOX genes (Kress et al., 2015), key developmental regulators of cell identity that are frequently deregulated in cancer (Yadav et al., 2024).

In the canonical model, MYC proteins heterodimerize with MAX, forming the transcriptionally active complex that regulates target gene expression, making MYC activity generally MAX-dependent (Dhanasekaran et al., 2022). Human cancers overexpress MYC factors at varying degrees and through distinct genetic mechanisms including chromosomal translocations, genomic amplification and promoter mutations (Dhanasekaran et al., 2022). A striking example is childhood neuroblastoma, in which MYCN is amplified to hundreds of copies (Schaub et al., 2018). This implies that upon MYC overexpression, MAX levels should not represent a limiting factor. However, large deletions or loss of function mutations in the MAX gene have been identified in a subset of tumors (Bausch et al., 2017), suggesting that MAX-independent functions of MYC also contribute to tumorigenesis. In line with that, the most aggressive neuroblastomas are those in which MYCN is overexpressed while MAX levels are below normal (Ferrucci et al., 2018). Recent work has also shown that MYCN modulates RNA metabolism upon recruiting the nuclear exosome, a 3’-5’ exoribonuclease complex, to its target genes and binding intronic transcripts, thus facilitating S-phase progression and enhancing stress tolerance of neuroblastoma cells (Papadopoulos et al., 2022), (Papadopoulos et al., 2024). These functions do not seem to directly require MAX and call for deeper investigation into how MYCN, and more broadly MYC proteins, influence cellular behavior and tumor progression outside of their dimerization with MAX.

To dissect oncogenic mechanisms controlled by MYC and/or MAX *in vivo*, it is essential to study the effects of altering their absolute and relative expression levels. The fruit fly, *Drosophila melanogaster*, provides an ideal system for this purpose, as it carries single orthologs (dMyc and dMax) and sophisticated genetic tools to constitutively or conditionally modulate gene expression are available. Many functions of the vertebrate MYC are conserved in flies (Grifoni & Bellosta, 2015), including regulation of cell growth (Secombe et al., 2007), (Steiger et al., 2008) and, during physiological cell competition, promotion of cell proliferation (Cova et al., 2004), (Moreno & Basler, 2004). Here, we demonstrate that a high dMyc/dMax ratio in undifferentiated eye progenitors disrupts cell differentiation and promotes eye-to-wing transdetermination by altering the physiological expression of HOX genes. Genetic and transcriptomic analyses reveal ectopic activation of the wing HOX gene *Antennapedia* and the wing developmental program. This is enhanced by loss of the HOX factor Deformed and accompanied by suppression of the eye developmental program. Our findings offer a novel perspective on MYC’s oncogenic role, highlighting cell fate reprogramming as a crucial mechanism through which MYC may influence tumor progression.

## Results

### Overexpressing fly dMyc or human MYCN in eye progenitors affects eye formation

To investigate the impact of high MYC levels on cell fate, we used the fly eye as an *in vivo* model. The eye develops from an eye primordium (imaginal disc), that forms in the embryo and proliferates in the larva. At late larval stage (wandering third larval instar, wL3), a wave of differentiation called morphogenetic furrow moves across the eye disc from posterior to anterior, promoting proliferation and differentiation of progenitors into photoreceptors. Photoreceptors organize in highly regular eye units, ommatidia, forming the adult compound eye (Baker, 2001), (Ready et al., 1976) (**Figure 1 A**). Using the UAS-Gal4 system (Brand & Perrimon, 1993), we overexpressed *Drosophila* MYC (dMyc) in undifferentiated eye imaginal disc progenitors by inducing a *UAS-dMyc* transgene with the *eyeless-Gal4* (*ey-Gal4*) driver (Halder et al., 1998), (Hauck et al., 1999). Expression of *ey-Gal4* was confirmed using a *UAS-dGFP* reporter, which encodes a rapid turnover (destabilized) GFP for sensitive detection of transient gene expression (X. Li et al., 1998) (**Supplementary Figure 1**). In these conditions, a twofold increase in dMyc transcripts is detected by qPCR analyses, while dMax (*Drosophila* Max) levels remain unchanged (**Supplementary Figure 2**). Notably, 75% of the animals show morphological defects, which can be grouped in two phenotypic classes based on the extent and intensity of the eye abnormalities: high-dMyc Class 1 - mild misalignment of the ommatidia confined in the posterior part of the eye (**Supplementary Figure 3**) or, in some cases, throughout the midline of the eye; high-dMyc Class 2 – cone-shaped eye (**Figure 1 B-D’**). To assess whether dMyc overexpression can affect eye development in a dose-dependent manner, we enhanced the activity of the UAS-Gal4 system by increasing the number of Gal4 and UAS transgenes carried by the flies and/or by raising the incubation temperature (mild activation at 24 °C; strong activation at 29 °C). Increasing the levels of dMyc overexpression indeed exacerbates the eye defects, as flies carrying one transgene copy (*1x ey-Gal4, 1x UAS-dMyc*) display more high-dMyc Class 1 and high-dMyc Class 2 eye defects when raised at 29°C (**Figure 1 E**). Flies with *2x ey-Gal4, 2x UAS-dMyc* could not be analyzed as they die at larval stage, likely due to *eyeless* expression in a few neuronal progenitors in the developing brain (Hauck et al., 1999), (Noveen et al., 2000). Importantly, similar eye phenotypes are observed upon overexpression of human MYCN (*UAS-MYCN*). Defects caused by either dMyc or MYCN overexpression can be rescued by the presence of a *dMyc* loss-of-function allele (*dMyc^G0354^*) (**Figure 1 E**)). This indicates functional conservation between fly and human MYC proteins.

**Figure 1.**
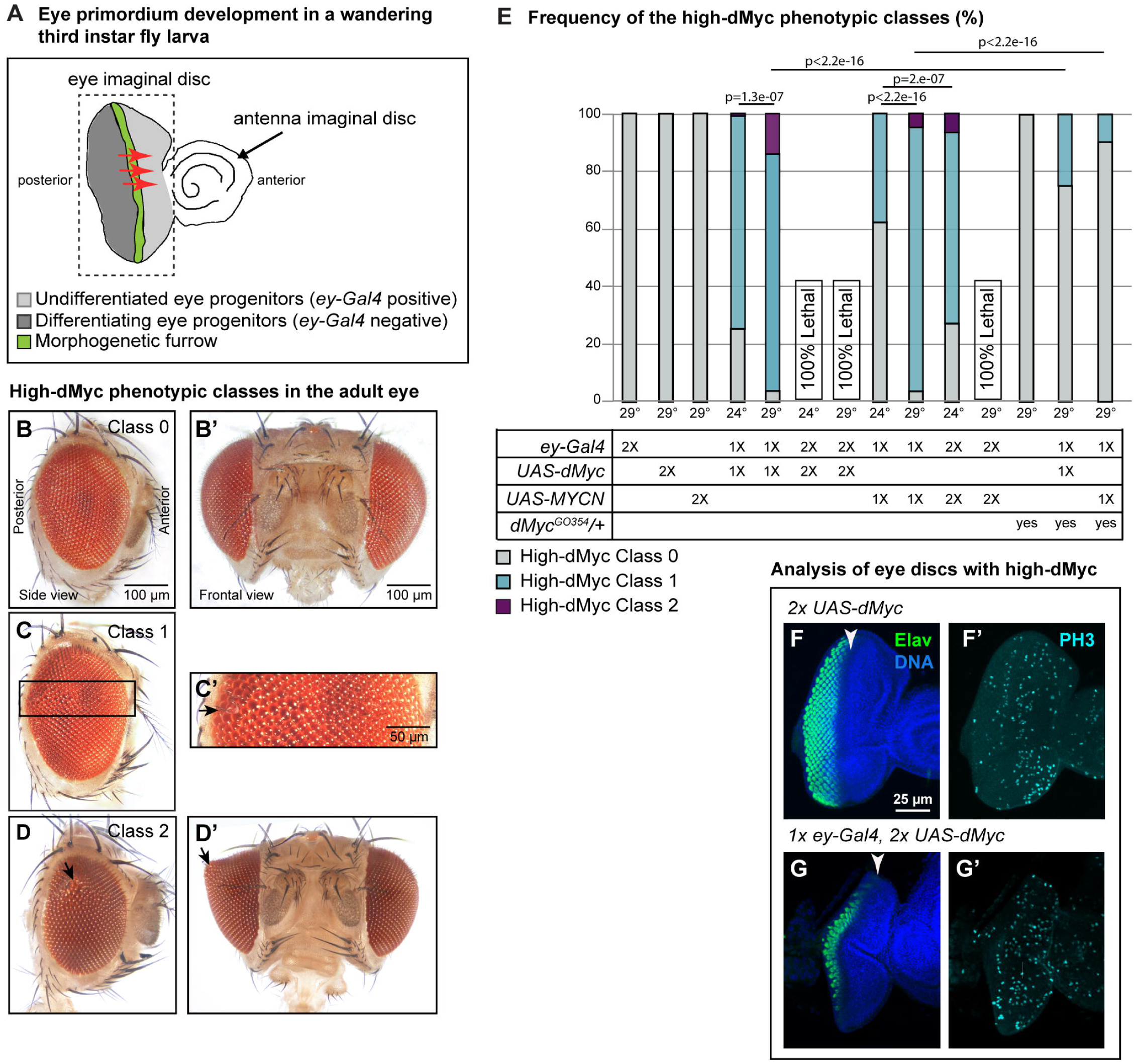
- Overexpression of *Drosophila* Myc or human MYCN in eye progenitors disrupts eye development. **(A)** Schematic of the eye development in a wandering third instar (wL3) fly larva. The *Drosophila* eye develops from the larval eye imaginal disc, which expands through progenitor proliferation. At the wL3 stage, the morphogenetic furrow sweeps from the posterior to the anterior eye disc margin (red arrows), triggering differentiation of photoreceptors, which will form the ∼800 ommatidia (eye units) that make up the adult compound eye. In the developing disc, only undifferentiated progenitors express the *ey-Gal4* driver, which is not active in differentiating photoreceptors posterior to the morphogenetic furrow **(B–D’)** Adult eye bright-field images showing the high-dMyc phenotypic classes observed upon overexpressing *Drosophila* Myc (*UAS-dMyc*) or the human MYCN (*UAS-MYCN*) in undifferentiated eye progenitors (*ey-Gal4*). Class 0: wild-type-like eyes **(B,B’)**; Class 1: mild ommatidial misalignment **(C,C’)**; Class 2: cone-shaped eyes. Arrows point to eye defects **(D,D’)**. The box in **(C)** indicates the magnified view in **(C’)**. **(E)** Quantification of the high-dMyc phenotypic classes. Genotypes (1X = one transgene copy, 2X = two transgene copies) and rearing temperature are indicated below the bar chart. Of note, both dMyc and MYCN overexpression phenotypes are rescued by heterozygous dMyc mutation (*dMyc^GO354^/+*). N≥150. Statistics: Mann-Whitney U test. **(F, G’)** Immunolabeling assay on eye discs from wL3 larvae upon dMyc overexpression in undifferentiated eye progenitors. Differentiating photoreceptors are labeled with anti-Elav (green), proliferating cells with anti-PH3 (cyan) and nuclei with Hoechst (blue). Arrowheads mark the position of the morphogenetic furrow.

Defective differentiation and arrangement of ommatidia in the adult eye can arise from alterations during eye disc development (Baker et al., 1990), (Baonza et al., 2001). To investigate this point, we performed immunolabeling assays on wL3 eye discs overexpressing dMyc. Although the eye discs appear smaller, cell proliferation, as assessed by anti-phosphorylated Histone H3 (PH3) labeling (Pérez-Cadahía et al., 2009), is not strikingly altered. By contrast, the number of cells expressing Elav, which is normally present in the differentiating photoreceptors posterior to the morphogenetic furrow (Robinow & White, 1991), (Robinow & White, 1988), is reduced, with in some cases only a few differentiating cells remaining in the disc (**Figure 1 F,G’**).

To summarize, dMyc overexpression impinges on eye cell differentiation, leading to morphological alterations of adult eyes.

### Fly dMax is required in undifferentiated eye progenitors to ensure cell proliferation and tissue growth

To explore whether dMax, like dMyc, contributes to cell differentiation control, we downregulated dMax in the undifferentiated eye progenitors using an RNAi construct targeting dMax (*UAS-dMax-RNAi*, hereafter *dMax-RNAi*) driven by *ey-Gal4*. This strategy significantly reduces the levels of dMax without affecting dMyc expression, as assessed by qPCR analyses (**Supplementary Figure 2**). Consistent with the known growth-promoting function of dMyc-dMax dimers in physiological and tumoral contexts, downregulation of dMax induces growth-defective eyes, while overall eye differentiation remains preserved (**Figure 2 A-C**). The morphological defects fall in two phenotypic classes of increasing severity: low-dMax Class 1 - showing mild loss of ommatidia; low-dMax Class 2 - characterized by a significantly reduced eye size (**Figure 2 A-C**). Introducing an extra copy of the *dMax* gene (*dMax-rescue* construct (Steiger et al., 2008)) rescues the eye phenotype in *1x ey-Gal4, 1x dMax-RNAi* animals. By contrast, enhancing UAS-Gal4 efficiency by increasing transgene copies or temperature worsens the eye phenotype and ultimately leads to lethality at the end of the pupal stage, as the imago (the adult fly) never emerges from the pupal case (*2x ey-Gal4, 2x dMax-RNAi* flies raised at 29°C) (**Figure 2 D**). These animals show small/absent eyes and a reduced head capsule (**Figure 2 E,F**), two structures both developing from the eye disc (Haynie & Bryant, 1986).

**Figure 2.**
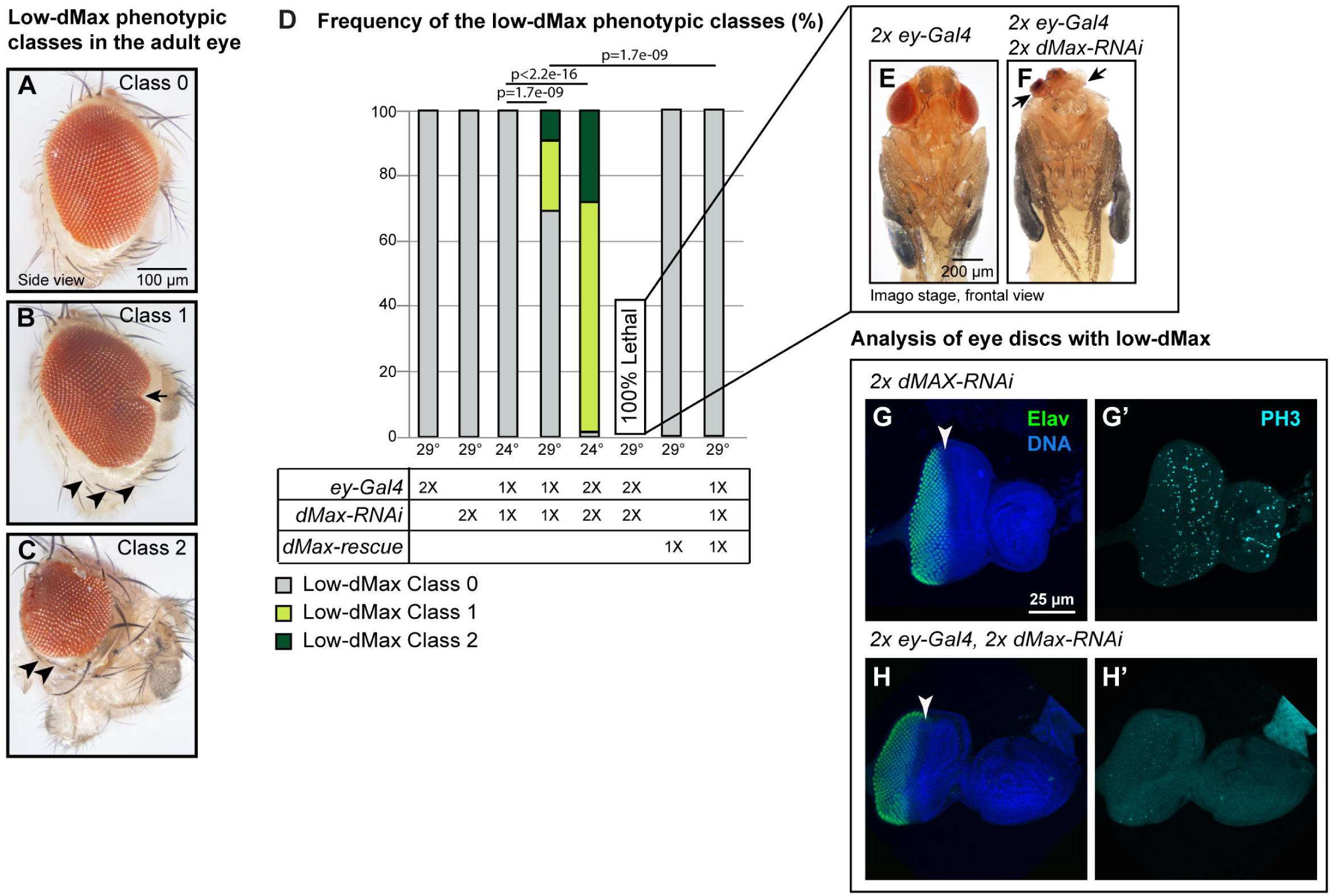
– Downregulation of *Drosophila* Max in eye progenitors reduces eye cell proliferation and growth. (A-C) Adult eye bright-field images showing the low-dMax phenotypic classes observed upon downregulating *Drosophila* Max (*dMax-RNAi*) in undifferentiated eye progenitors (*ey-Gal4*). Class 0: wild-type-like eyes **(A)**; Class 1: mild growth defect **(B)**; Class 2: small eye **(C)**. Arrows point to eye defects. Arrowheads indicate periorbital vibrissae, bristles that mark the perimeter of the adult eye territory. **(D)** Quantification of the low-dMax phenotypic classes. Genotypes (1X = one transgene copy, 2X = two transgene copies) and rearing temperature are indicated below the bar chart. Of note, dMax downregulation phenotype is rescued by an extra copy of dMax (*dMax-rescue*). N≥130. Statistics: Mann-Whitney U test. **(E, F)** Bright-field images showing flies at imago stage upon dMax downregulation in undifferentiated eye progenitors. Arrows point to eye and head defects. **(G-H’)** Immunolabeling assay on eye discs from wL3 larvae upon dMax downregulation in undifferentiated eye progenitors. Differentiating photoreceptors are labeled with anti-Elav (green), proliferating cells with anti-PH3 (cyan) and nuclei with Hoechst (blue). Arrowheads mark the position of the morphogenetic furrow.

A small eye disc induced by dMax downregulation could therefore account for the size-reduction in the adult eye and, in the most severe cases, adult head. Immunolabeling analyses on eye discs collected from larvae upon dMax downregulation confirm this hypothesis. Eye discs are smaller, and proliferation is strongly reduced throughout the disc (**Figure 2 G,H’**). Unlike upon dMyc overexpression (**Figure 1 F,G’**), cell differentiation is maintained upon dMax silencing, as both the progression of the morphogenetic furrow and the appearance of Elav+ differentiating photoreceptors are unaffected (**Figure 2 G,H’**).

Altogether, these findings indicate that dMax promotes cell proliferation and tissue growth in the developing eye, yet it is dispensable for progenitor differentiation.

### Increasing dMyc/dMax ratio triggers cell transdetermination

Our data suggest that dMyc regulates cell differentiation independently of dMax, as high dMyc expression, but not dMax downregulation, impairs Elav expression (the classical marker of eye progenitor differentiation). If this is correct, increasing the dMyc/dMax intracellular ratio by simultaneously combining dMyc overexpression and dMax downregulation should exacerbate the differentiation defects. Indeed, lowering dMax levels would reduce the pool of dMyc-dMax dimers and increase the dMyc not engaged in dimerization, hence allowing the dMax-independent activity of dMyc. To test this hypothesis, we generated flies carrying *ey-Gal4*, *UAS-dMyc* and *Max-RNAi* transgenes. While 1*x ey-Gal4*, *1x UAS-dMyc*, *1x dMax-RNAi* flies raised at 24°C are similar to controls, one third of those raised at 29°C exhibit a spectrum of mild phenotypes (mild high-dMyc-low-dMax Class 1, **Figure 3 A-B’**), recapitulating the ommatidial loss (**Figure 3 B’** asterisks) observed upon dMax downregulation (**Figure 2 A-C**), and ommatidia misalignment (**Figure 3 B’** arrows) observed upon dMyc overexpression (**Figure 1 B-D’**), but not in dMax-silenced eyes (please compare **Figure 2 B** and **Figure 3 B,B’**).

**Figure 3.**
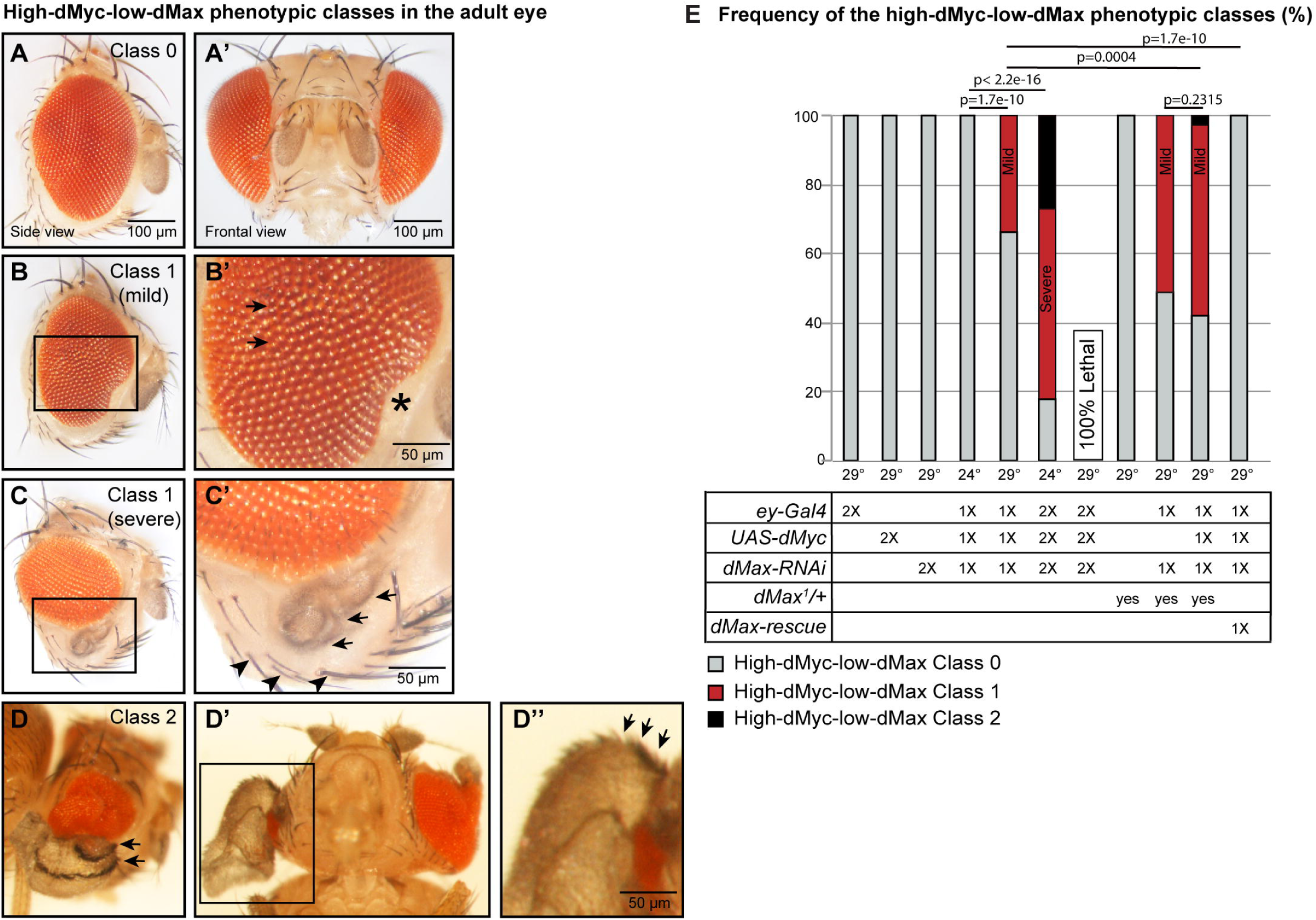
– Enhancing dMyc/dMax ratio in eye progenitors induces cell transdetermination. (A-D”) Adult eye bright-field images showing the high-dMyc-low-dMax phenotypic classes observed upon overexpressing dMyc (*UAS-dMyc*) and downregulating dMax (*dMax-RNAi*) in undifferentiated eye progenitors (*ey-Gal4*), as well as upon strong but viable dMax downregulation (*dMax-RNAi* driven by *ey-Gal4* in animals carrying heterozygous *dMax^1^* mutation). Class 0: wild-type-like eyes **(A,A’)**; Class 1: phenotypic spectrum ranging from mild cases in which some ommatidia are missing (asterisk) and others fail to properly align in the compound eye (arrows) **(B,B’)**, to severe cases in which ectopic, bristle-decorated tissue (arrows) develops in the eye territory delimited by periorbital vibrissae (arrowheads) **(C,C’)**; Class 2: wing-like structure in the eye territory characterized by bristles resembling those associated to the wing (arrows) **(D-D”)**. **(E)** Quantification of the high-dMyc-low-dMax phenotypic classes. Genotypes (1x = one transgene copy, 2x = two transgene copies) and rearing temperature are indicated below the bar chart. *2x ey-Gal4*, *2x UAS-dMyc*, *2x dMax-RNAi* die when raised at 29°C (likely due to dMyc overexpression in few eyeless-expressing brain neuroblasts, similarly to *2x ey-Gal4*, *2x UAS-dMyc.* Of note, high-dMyc-low-dMax phenotype is rescued by an extra copy of dMax (*dMax-rescue*). N≥200. Statistics: Mann-Whitney U test.

Importantly, increasing the number of transgene copies worsens the ommatidia phenotypes (**Figure 3 E**). Indeed, 82% of *2x ey-Gal4*, *2x UAS-dMyc*, *2x dMax-RNAi* flies raised at 24°C exhibit eye defects. 55% of those flies corresponds to severe high-dMyc-low-dMax Class 1 phenotypes (**Figure 3 C,C’**) in which ventral ommatidia are replaced by ectopic, bristle-decorated tissue (**Figure 3 C’** arrows). This tissue still belongs to the eye territory as it lies internal to the periorbital vibrissae (**Figure 3 C’** arrowheads), bristles that mark the perimeter of the adult eye territory (Ouweneel, 1970), (Haynie & Bryant, 1986). This suggests that part of the eye fails to differentiate into ommatidia and instead gives rise to other cell types, in contrast to the low-dMax Class 2 phenotype (**Figure 2C**), where ommatidia are tightly surrounded by vibrissae, indicating a smaller but fully differentiated eye. Moreover, in 27% of cases, we observe a wing-like structure within the eye territory (high-dMyc-low-dMax Class 2, **Figure 3 D-E**). The wing-like structure displays varying degrees of organization, ranging from a wing bud to a clearly recognizable, unfolded wing (**Figure 3 D-D’’**). In the latter case, they also contain sensory bristles characteristic of the anterior wing margin (Hartenstein & Posakony, 1989) (**Figure 3 D,D’’** arrows). This could be assimilated to those observed in severe high-dMyc-low-dMax Class 1 flies (**Figure 3C’** arrows), suggesting phenotypic continuity between high-dMyc-low-dMax Class 1 and high-dMyc-low-dMax Class 2.

The development of a wing-like structure in the eye represents a typical case of cell transdetermination, a process in which a cell switches fate and differentiates into another cell type.

Importantly, no ectopic wing-like structures are observed when dMyc overexpression nor dMax downregulation are carried out separately, suggesting that the dMyc/dMax intracellular ratio more than their absolute levels accounts for dMyc-mediated impact on cell fate and differentiation. To formally prove this point, we generated flies with low, yet viable dMax levels by downregulating dMax in heterozygous *dMax* mutants (*1x ey-Gal4, 1x dMax-RNAi, dMax^1^/+*). Strikingly, 51% of these flies show mild high-dMyc-lox-dMax Class 1 phenotypes. Further increase of the dMyc/dMax ratio by adding a *UAS-dMyc* transgene (*1x ey-Gal41, 1x UAS-dMyc, 1x dMax-RNAi, dMax^1^/+*) leads to the appearance of high-dMyc-low-dMax Class 2 eyes and exacerbates the eye defects observed in *1x ey-Gal41, 1x UAS-dMyc, 1x dMax-RNAi* animals. Conversely, lowering the dMyc/dMax ratio by introducing an extra copy of the *dMax* gene (*1x ey-Gal41, 1x UAS-dMyc, 1x dMax-RNAi, dMax-rescue*) rescues the eye phenotype (**Figure 3 E**).

Take together, these data demonstrate that the relative intracellular levels of dMyc and dMax regulate cell differentiation, with a high dMyc/dMax ratio triggering eye-to-wing transdetermination.

### The physiological HO X program is affected by high dMyc/dMax ratio

Cell identity is tightly regulated by HOX genes, evolutionarily conserved transcription factors whose misexpression is known to produce homeotic transformations, where one body part is converted into another (Emerald & Roy, 1997). The HOX gene *Antennapedia (Antp)* is normally expressed in the wing disc (Levine et al., 1983), where it antagonizes eye selector genes, thus inhibiting ectopic eye development (Plaza et al., 2001). Ectopic Antp in the eye disc leads to an eye-to-wing transformation (Kurata et al., 2000), (Prince et al., 2008).

Given the above-mentioned phenotypes, we asked whether increasing dMyc/dMax ratio in eye progenitors results in ectopic expression of Antp. Remarkably, immunolabeling assays on eye disc of *2x ey-Gal4, 2x UAS-dMyc, 2x dMax-RNAi* wL3 larvae reveal the presence of Antp-expressing cells, which in extreme cases cluster together and associate with severe morphological defects of the eye disc (**Figure 4 A-B’’**). Few Antp+ cells also express the neuronal marker Elav (**Supplementary Figure 4 A-B’’**), yet none of them express the mitotic marker PH3 (**Figure 4 B’’, Supplementary Figure 4 C-D’’**), suggesting that they have exited the cell cycle possibly to start differentiating. Importantly, Antp is also detected upon dMyc, as well as MYCN, overexpression or strong dMax downregulation (**Supplementary Figure 5 A-D”’**). Yet, in these cases Antp-expressing cells are scattered and positive for PH3, and the eye disc morphology is preserved (**Supplementary Figure 5 A-D”’**). Consistently, these combinations do not lead to a wing-like structure in the eye territory (**Figure 1 B-D’** and **Figure 3 A-E**). Together, these findings suggest that an increased dMyc/dMax ratio is sufficient to induce ectopic Antp expression in the eye disc and, in severe cases, activate a wing fate program.

**Figure 4.**
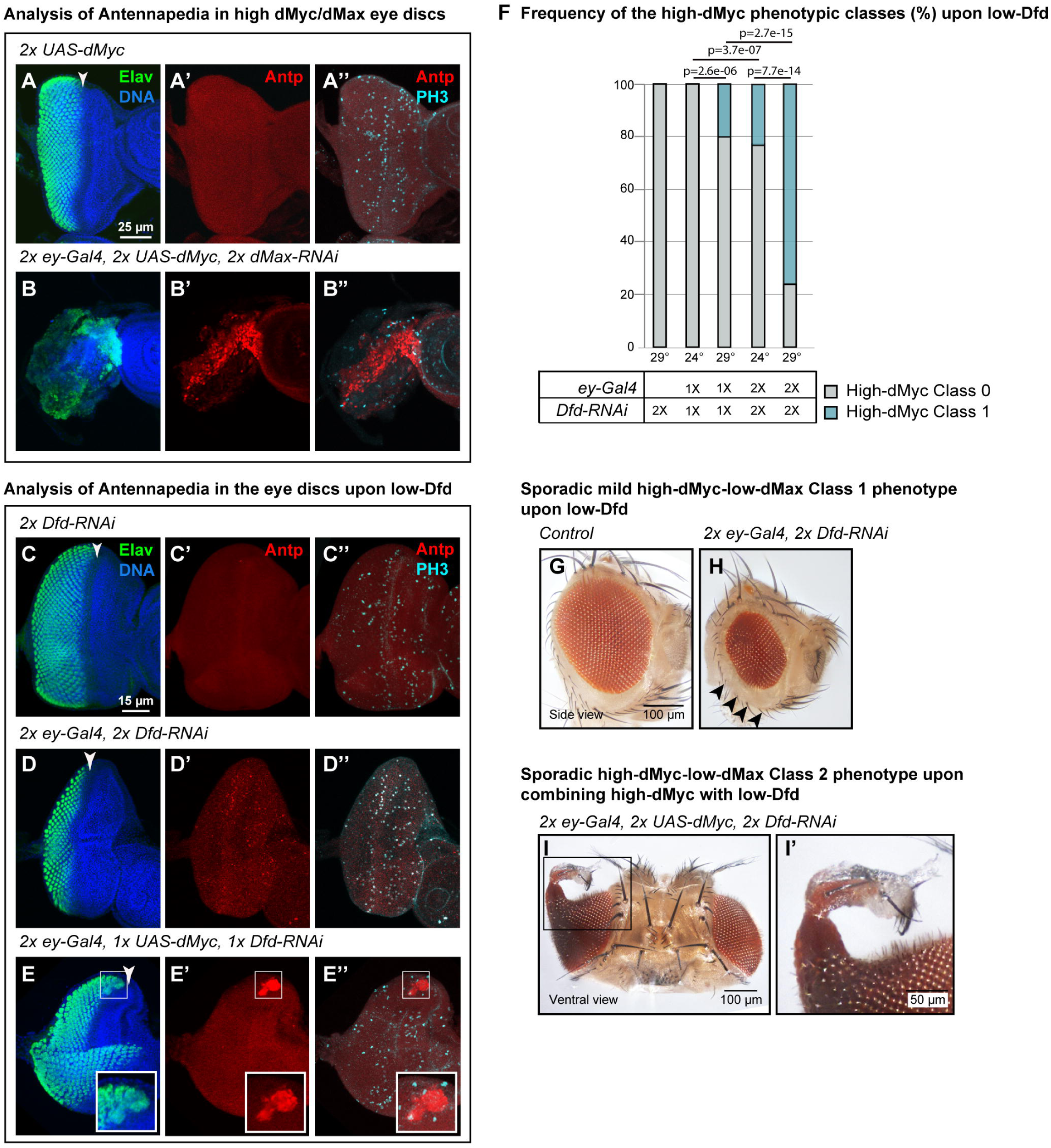
– High dMyc/dMax ratio impacts the eye HOX program. (A-E”) Immunolabeling assay on eye discs from wL3 larvae upon **(A-B”)** dMyc overexpression and dMax downregulation, **(C-D”)** downregulation of Deformed (*Dfd-RNAi*), **(E-E”)** dMyc overexpression and Dfd downregulation in undifferentiated eye progenitors (*ey-Gal4*). Differentiating photoreceptors are labeled with anti-Elav (green), Antennapedia positive cells with anti-Antp (red), proliferating cells with anti-PH3 (cyan) and nuclei with Hoechst (blue). Arrowheads mark the position of the morphogenetic furrow. **(F)** Quantification of the high-dMyc phenotypic classes observed upon Dfd downregulation. Genotypes (1X = one transgene copy, 2X = two transgene copies) and rearing temperature are indicated below the bar chart. N≥200. Statistics: Mann-Whitney U test. **(G,H)** Adult eye bright-field images showing mild high-dMyc-low-dMax Class 1 phenotype occasionally observed upon Dfd downregulation in undifferentiated eye progenitors **(H)** compared to a wild-type like eye of the same genotype **(G)**. Arrowheads indicate periorbital vibrissae. **(I, I’)** Adult eye bright-field images showing a wing-like structure (high-dMyc-low-dMax Class 2 phenotype) occasionally observed upon combining Dfd downregulation with dMyc overexpression in undifferentiated eye progenitors.

Gain- or loss-of-function mutations in a Hox gene can alter segment identity by changing where that Hox gene itself is expressed and by disrupting the normal repression/activation interactions among neighboring Hox genes (Mallo & Alonso, 2013). The HOX gene *Deformed (Dfd)* is expressed in the eye disc (Diederich et al., 1991), (Chadwick et al., 1990), (Martinez-Arias et al., 1987), and *Dfd* loss-of-function mutations are known to result in transformations towards thoracic structures (Merrill et al., 1987), (Regulski et al., 1987), (Hughes & Kaufman, 2002). Being the wing a thoracic appendage, we speculated that ectopic Antp expression in the eye disc upon high dMyc/dMax ratio may be epistatic to Dfd. Dfd silencing in eye progenitors (*ey-Gal4, Dfd-RNAi*) is sufficient to drive ectopic expression of Antp in proliferating cells of the eye disc (**Figure 4 C-D’’**) and mimics dMyc overexpression phenotypes in the adult eye in a dose-dependent fashion (**Figure 4 F**). Interestingly, eye defects resembling mild high dMyc-low-dMax Class 1 (**Figure 3 B-C’**) are also occasionally observed upon Dfd downregulation (**Figure 4 G,H**), and they become more severe upon further dMyc overexpression, supporting a genetic interaction between Dfd silencing and dMyc overexpression. Indeed, *2x ey-Gal4*, *2x UAS-dMyc, 2x Dfd-RNAi* flies in rare cases exhibit a tissue overgrowth bulging from the eye territory (**Figure 4 I,I’**), resembling a rudimental wing-like structure similarly to what is induced by high dMyc/dMax ratio (**Figure 3 D-D’’**). Accordingly, the eye discs with Dfd downregulation and dMyc overexpression occasionally show clustered, non-proliferating Antp+ cells and disorganized Elav labeling (**Figure 4 E-E’’**), reminiscent of the phenotypes observed under high dMyc/dMax ratio (**Figure 4 B-B’’**).

Overall, these data demonstrate that loss of Dfd enables ectopic expression of Antp and synergizes with elevated dMyc levels to induce cell transdetermination, revealing HOX genes as key mediators of MYC-driven regulation of cell identity.

### High dMyc/dMax ratio globally disrupts the developing program of the eye progenitors

To gain a genome-wide and unbiased perspective on the molecular consequences of a high dMyc/dMax ratio in undifferentiated eye progenitors, we performed a bulk RNA sequencing (RNA-seq) assay on eye imaginal discs collected from *2x ey-Gal4, 2x UAS-dMyc, 2x dMax-RNAi* wL3 larvae raised at 24°C or, as a control, from larvae with equal number of UAS and Gal4 transgenes (*2x ey-Gal4, 4x GFP-RNAi*). For each genotype, we performed three independent replicates, which group together in a Principal Component Analysis (**Supplementary Figure 6**) indicating consistency across replicates. Among the detected genes (**Supplementary Table 1**), we defined differentially expressed genes (DEGs) as those with |log2FC(high-dMyc-low-dMax/control)|>1, adjusted p-value < 0.05. A baseMean ≥ 14 (25th percentile cutoff) was used to exclude lowly expressed genes.

Out of the 835 DEGs identified, 314 genes are upregulated and 521 are downregulated (**Supplementary Table 1**). Interestingly, and in agreement with the observed phenotypes, gene ontology (GO) analysis of the downregulated genes highlights several eye development GO terms (**Figure 5 A, Supplementary Table 2**). These include key eye regulators such as the Pax-family transcription factor Eyegone (Eyg), that promotes eye disc proliferation (Jang et al., 2003); the transcription factor Prospero (Pros) expressed in R7 photoreceptors (Kauffmann et al., 1996) and required for their differentiation (Cook et al., 2003); the transcription factor Retained (Retn) (Ditch et al., 2005), and the Heat shock protein 27 (Hsp27), whose downregulation affects ommatidia morphology producing rough eye phenotypes (Chen et al., 2012). Moreover, the RNA-binding protein Embryonic lethal abnormal vision (Elav) expressed in differentiating photoreceptors (Robinow & White, 1988) and whose mutation produce aberrant eye structures (Homyk et al., 1985) also shows a tendency to decrease (log2FC=-0.7) (**Figure 5 B**). Overall, this indicates that a global suppression of the eye developmental program contributes to the morphological defects observed in adult eyes (**Figure 3 A–D’’**).

**Figure 5.**
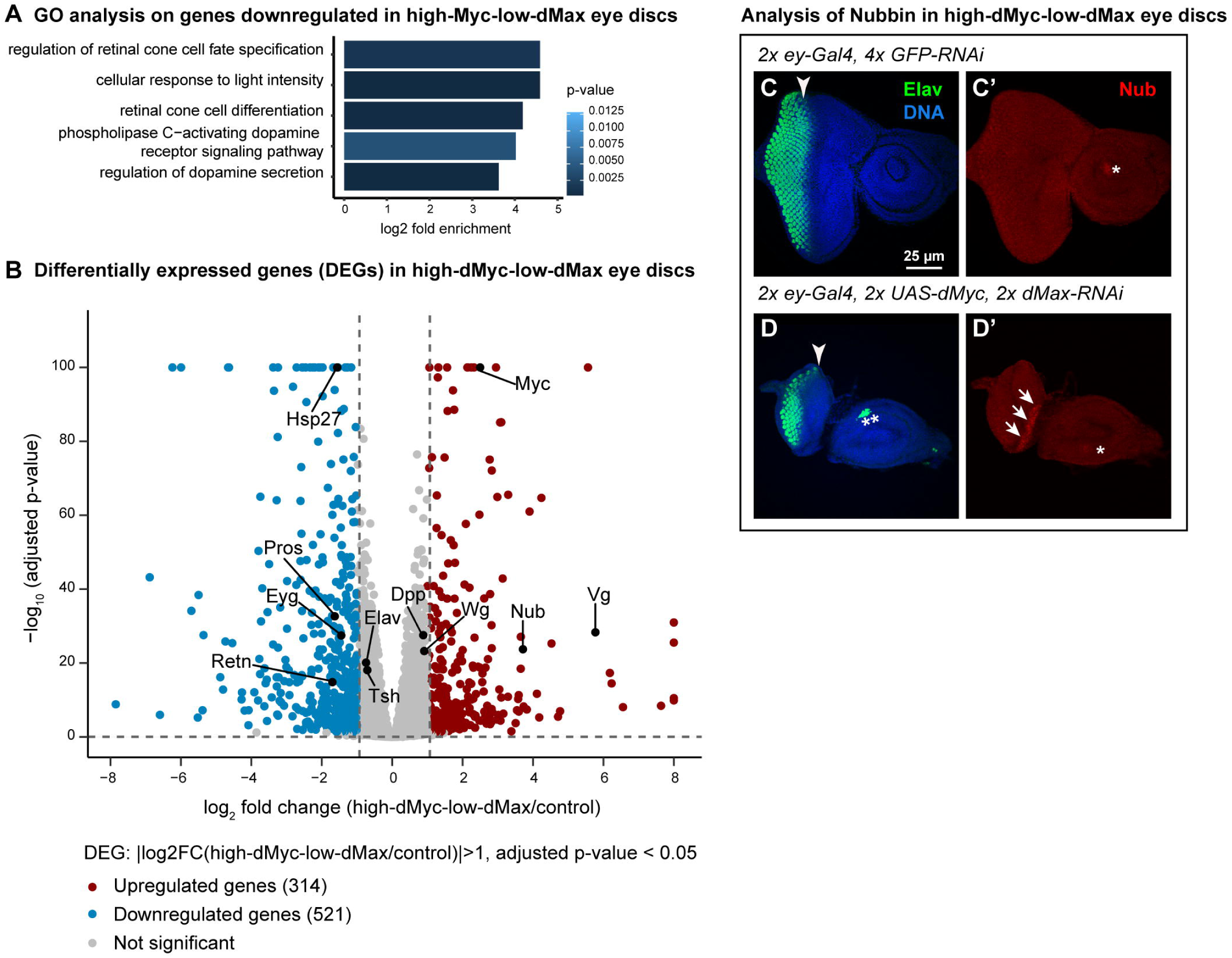
– High dMyc/dMax ratio induces transcriptome-wide changes in the developing eye. **(A)** Gene ontology (GO) analysis on the downregulated genes identified in the bulk RNA-seq assay on high-dMyc-low-dMax (*2x ey-Gal4, 2x UAS-dMyc, 2x dMax-RNAi*) and control (*2x ey-Gal4, 4x GFP-RNAi*) wL3 eye discs (∼120 hour after egg lay at 24°C). Differentially expressed genes (DEGs) were defined based on |log2FC(high-dMyc-low-dMax/control)|>1, adjusted p-value < 0.05 and baseMean ≥ 14 (25th percentile cutoff). **(B)** Volcano plot of bulk RNA sequencing analysis. Each dot represents a gene. The 314 upregulated genes are shown in red, the 521 downregulated genes are shown in blue. Selected genes with roles in eye and wing development are labeled. **(C-D’)** Immunolabeling assay on eye discs from wL3 larvae upon dMyc overexpression and dMax downregulation in undifferentiated eye progenitors. Differentiating photoreceptors are labeled with anti-Elav (green), Nubbin positive cells with anti-Nub (red) and nuclei with Hoechst (blue). Arrowheads mark the position of the morphogenetic furrow. Arrows point to ectopic expression of Nub in the eye disc, whereas asterisks mark expected Nub expression in the antenna disc. Artifactual Elav expression in **(D)** is indicated by a double asterisk.

Importantly, wing development genes are upregulated in high-dMyc-low-dMax eye discs, including the transcription factors Vestigial (Vg), essential for wing identity as its mutation results in significantly reduced wings (Williams & Bell, 1988), and Nubbin (Nub), expressed in the nascent wing and required for proper proximo-distal wing patterning (Ng et al., 1995) (**Figure 5 B**). Two major interactors of Vg also show a tendency to increase (log2FC=0.9): the TGFβ ligand Decapentaplegic (Dpp) (Mohit et al., 2006) and the WNT factor Wingless (Wg) (Klein & Arias, 1998), (Baena-López & García-Bellido, 2003) (**Figure 5 B**), which create antero-posterior and dorso-ventral gradients essential for wing development and sensory organ formation (Neumann & Cohen, 1996), (Nellen et al., 1996). Together with the proximo-distal information from Nub, these signals may provide a molecular basis for wing bud polarization, ultimately explaining the emergence of a wing-like structure with sensory bristles from the eye territory (**Figure 3 D-D’’**). Immunolabeling assays validate the ectopic Nub expression in high-dMyc-low-dMax eye discs (example shown in **Figure 5 C’,D’**), as detected by our RNA-seq analysis.

In summary, a high dMyc/dMax ratio in undifferentiated eye progenitors impacts the HOX program and broadly suppresses the eye developmental cascade while ectopically activating a wing fate program.

## Discussion

The MYC family of transcription factors are well-known oncoproteins that, through dimerization with MAX, regulate key processes such as cell growth, proliferation and differentiation. However, in many cancers, MYC/MAX ratio is increased by MYC overexpression or MAX reduction (Dhanasekaran et al., 2022), (Ferrucci et al., 2018), (Bausch et al., 2017), (Burnichon et al., 2012), (Comino-Méndez et al., 2011), raising the possibility of a MAX-independent role of MYC. Using the *Drosophila* eye as an *in vivo* model, we identify MYC/MAX stoichiometry as a critical regulator of cell identity and differentiation. Elevated dMyc levels relative to dMax induces cell transdetermination culminating in an eye-to wing homeotic transformation. These findings reveal a MAX-independent role of dMyc in controlling cell identity, with broad implications for developmental biology and MYC-driven tumors.

### High dMyc/Max ratio drives cell transdetermination while inducing a pro-oncogenic signature

Consistent with the role of dMyc–dMax dimers in promoting proliferation, dMax downregulation in undifferentiated eye progenitors strongly reduces proliferation within the developing eye (eye imaginal disc), resulting in a small adult eye. By contrast, dMyc overexpression does not enhance progenitor proliferation. This is likely not due to limiting dMax, as a similar approach of dMyc overexpression using the *GMR-Gal4* driver (expressed in differentiating eye cells) does increase cell size in the adult eye (Secombe et al., 2007) in a Max-dependent way (Steiger et al., 2008), indicating that dMax levels are sufficient for dMyc–dMax function. Instead, excess dMyc in undifferentiated eye progenitors interferes with cell differentiation, producing cone-shaped eyes (high-dMyc Class 2), suggesting that dMyc functions in a context-dependent manner depending on whether it is overexpressed in undifferentiated cells or in cells already committed to terminal differentiation. Increasing dMax expression rescues the differentiation defects (**Supplementary Figure 7**), demonstrating that they arise from enhanced dMyc/dMax ratio, rather than absolute dMyc levels. Moreover, further raising this ratio by combining dMyc overexpression with dMax silencing aggravates differentiation defects and induces transdetermination, converting eye progenitors into a wing-like structure.

Intriguingly, gene ontology (GO) analysis of genes upregulated in high-dMyc/low-dMax discs revealed enrichment of terms related to dMyc’s oncogenic functions. This includes biological processes, such as trehalose metabolism, L-ornithine transport, and acetaldehyde metabolism (**Supplementary Figure 8**). Trehalose regulates organ size in flies by buffering glucose (Yasugi et al., 2017), and its levels increase in fly tumor models where dMyc is upregulated (Prober & Edgar, 2002), (Khezri et al., 2021). L-ornithine transport fuels polyamine synthesis, essential for cell proliferation in flies (Callaerts et al., 1992), (Burnette & Zartman, 2015) and in mammals (Sánchez-Jiménez et al., 2019), as well as growth of MYC-associated tumors (Nilsson et al., 2005), including neuroblastoma (Gamble et al., 2021). Aldehyde dehydrogenases, key enzymes in the acetaldehyde metabolism, promote synthesis of polyamines and mark stem cell populations in several human cancers (Singh et al., 2013), (Ginestier et al., 2007), (Feldmann et al., 2007). This suggest that, although we did not observe overt tumoral growth in eye disc with elevated dMyc/dMax ratio, the pro-oncogenic cascade driven by high dMyc may already begin to be activated.

Together, these findings indicate that dMyc/dMax stoichiometry governs cell growth and differentiation in a context-dependent manner, with a high dMyc/dMax ratio acting as a developmental switch that reprograms the fate of undifferentiated progenitors, while priming cells for tumorigenic transformation.

### High dMyc/dMax ratio impacts developmental programs and HOX gene expression

At the molecular level, the observed transdetermination resembles classical homeotic transformations and aligns with the role of HOX genes in maintaining segmental identity. Immunolabeling analyses revealed ectopic expression of the thoracic HOX gene *Antennapedia* (*Antp*) in high-dMyc-low-dMax eye discs, ranging from isolated Antp-expressing cells to Antp-positive clusters within morphologically altered tissue (**Figure 4 B,B’’**). This variability is consistent with the 27% penetrance of eye-to-wing transformation and likely caused Antp transcripts to fall below the detection threshold of bulk RNA sequencing. Nevertheless, our transcriptomic analysis identified a dual mechanism under high-dMyc-low-dMax conditions: (1) repression of eye differentiation genes (*eyegone, prospero, elav*), and (2) activation of wing determinants such as *nubbin* and *vestigial*, the latter being previously shown to be induced in the eye disc upon Antp overexpression (Kurata et al., 2000).

Multiple mechanisms may drive ectopic Antp expression. RNA sequencing data revealed downregulation of *teashirt* (*tsh*) (log2FC = –0.7, **Figure 5A**), a zinc finger transcription factor that cooperates with HOX proteins to define head identity by repressing Antp and whose loss induces ectopic Antp expression (Bhojwani et al., 1997). Because HOX domains are stabilized through mutual repression, we hypothesize that a dMyc/dMax ratio brought to the extreme also reduces expression of cephalic HOX genes such as *Deformed* (*Dfd*), which normally inhibits thoracic gene activity (Merrill et al., 1987), (Regulski et al., 1987), (Hughes & Kaufman, 2002). In agreement with this hypothesis, loss of Dfd phenocopies high-dMyc-low-dMax eye defects and synergizes with elevated dMyc to enhance Antp activation and cell transdetermination, indicating genetic crosstalk between these HOX genes.

Of note, besides the well-recognized transcriptional activation role of MYC, which is MAX-dependent, MYC has also been shown to repress transcription, and this function does not necessarily require MYC–MAX dimers (Brenner et al., 2005), (Gartel et al., 2001). Typically, in neuroblastoma, MYCN represses the TRKA and p75NTR neurotrophin receptors, whose robust expression predicts favorable prognosis, by binding their promoters and recruiting the histone deacetylase 1 (HDAC1) to establish a repressive chromatin state (Iraci et al., 2011). MYCN also recruit HDAC1 to represses tissue transglutaminase (TG2) and inhibit neuroblastoma cell differentiation (Liu et al., 2007). Similarly, high dMyc may act on chromatin to repress Dfd, thus inducing ectopic Antp expression and cell transdetermination. In support of this model, the eye phenotypes observed upon dMyc overexpression are partially rescued by heterozygous null mutations of dHDAC1 (*HDAC1^04556/+^*) or the *Drosophila* homolog of human NCoR/SMRT, Smrter (*Smr^G0060/+^*), that stabilizes HDAC recruitment into transcriptional repressor complexes (Mottis et al., 2013) (**Supplementary Figure 9**). The downregulation of dHDAC1 or Smrter *via* RNAi in undifferentiated eye progenitors confirmed these results (**Supplementary Figure 9**).

Collectively, these findings reveal that dMyc/dMax stoichiometry contributes to regulating the HOX network possibly through chromatin-repressive mechanisms, thereby safeguarding cellular identity.

### Pathological implications: a cancer perspective

Across the 33 cancer types in The Cancer Genome Atlas (TCGA), about 28% of samples show amplification of a MYC gene (Schaub et al., 2018). Our findings in the *Drosophila* model suggest that this amplification may not only sustain uncontrolled growth but also trigger cell fate reprogramming. Fate reprogramming is particularly relevant in cancer cells, which can acquire or lose features, thereby increasing their heterogeneity and adapting to changing microenvironments. This can promote tumor growth and dissemination, enhance treatment resistance, and crucially influence clinical behavior (Scully et al., 2012), (Fedele et al., 2024).

MYC factors are evolutionarily conserved structurally but also functionally, as overexpression of human MYCN reproduces the phenotypes induced by the *Drosophila* dMyc, including ectopic activation of the wing HOX gene *Antp* in the eye disc, phenotypes rescued by *dMyc* mutation. Once confirmed in humans, our model may help expand the known molecular bases of MYC oncogenic functions and help elucidating why MYC-driven tumors adopt hybrid differentiation states. Rather than solely inhibiting MYC, restoring proper MYC/MAX balance could redirect tumor cells toward a less aggressive, more differentiated state.

Aberrant dysregulation of HOX genes in cancer has been described in several studies. MYC is known to reshape chromatin architecture and globally amplify transcription, enabling inappropriate expression of lineage-restricted *loci* such as HOX clusters (Kress et al., 2015). In hematologic malignancies, MYC overexpression correlates with upregulation of *HOXA9* and *HOXA10*, genes normally required for early hematopoietic patterning but strongly leukemogenic when ectopically expressed (B. Li et al., 2019). Mechanistically, MYC recruits histone acetyltransferases and other chromatin modifiers to developmental loci, increasing enhancer accessibility and promoting activation of HOX-driven transcriptional programs (Schaub et al., 2018). Polycomb repression, which typically silences HOX clusters in differentiated tissues, is also disrupted by MYC-mediated epigenetic remodeling, further facilitating HOX de-repression (Parreno et al., 2022).

Functional interactions between MYC and HOX proteins reinforce these oncogenic effects. In acute myeloid leukemia, HOXA9 and MYC co-occupy enhancers governing proliferation and survival genes, and HOXA9 requires MYC to maintain leukemic stem cell identity (Miyamoto et al., 2021). Conversely, MYC-driven lymphomas exhibit widespread dysregulation of HOX family members, which contributes to blocked differentiation and enhanced tumor aggressiveness (Pelengaris & Khan, 2003).

In conclusion, unbalancing the MYC/MAX ratio may generate another layer of control to the MYC–HOX axis forming a feed-forward oncogenic circuit, impacting on tumor de-differentiation, cell-fate reprogramming, survival, and progression.

### Evolutionary implications

Beyond its developmental and pathological relevance, our work raises the intriguing possibility that MYC/MAX stoichiometry may have played a fundamental role in the evolution of metazoan body plans and lineage diversification. Both MYC and MAX are deeply conserved across animals, and their partnership is essential for coordinating growth, metabolic activity, and transcriptional output (Young et al., 2011). Our findings suggest that fluctuations in MYC/MAX balance can modulate HOX gene expression and thereby influence segmental identity, consistent with evidence that HOX boundaries and axial patterning respond to growth and metabolic regulators (Soshnikova & Duboule, 2009). Because HOX genes orchestrate the regionalization of the body axis, changes in MYC/MAX ratio could theoretically bias developmental programs toward alternative morphological outcomes, a concept aligned with evo-devo studies showing that small shifts in HOX regulation drive morphological diversification (Carroll, 2008). Such a mechanism may have contributed to the evolutionary plasticity of appendages and organs, especially in lineages where MYC dosage or MAX availability became rewired through gene duplication, promoter evolution, or chromatin changes (Soshnikova & Duboule, 2009). The sensitivity of HOX boundaries to MYC/MAX balance may have provided a means to fine-tune segmental identity or generate novel structures, offering a potential route by which metabolic regulators, growth factors, and patterning cues co-evolved (Hong et al., 2016). Thus, MYC/MAX stoichiometry may emerge not only as a determinant of cell fate but also as a potential evolutionary lever shaping morphological diversity among MYC/MAX-expressing organisms.

## Conclusion

In conclusion, this study discloses the MYC/MAX stoichiometry as a critical regulator of cell fate reprogramming, extending MYC’s role beyond mere growth control to the maintenance of cellular identity. This mechanism may contribute to the well-known cellular heterogeneity of MYC-driven cancers, which is difficult to study *in vivo* and may underlie therapeutic resistance.

### Limitations of the study

Despite the genetic and molecular evidence supporting an interaction between an elevated dMyc/dMax ratio and *Deformed* regulation, we were unable to directly detect a reduction of *Dfd* expression in high-dMyc-low-dMax eye discs, due to biological and technical constraints. In particular, the available anti-Deformed antibody provides weak and unreliable staining in imaginal discs, preventing a confident assessment of changes in protein levels. Moreover, *Dfd* is expressed in a very restricted domain of the eye disc (Diederich et al., 1991), (Chadwick et al., 1990), (Martinez-Arias et al., 1987), making detection challenging and placing it below the sensitivity threshold of bulk RNA-seq. Finally, the incomplete penetrance and variable expressivity of the high-dMyc-low-dMax phenotype hindered our ability to quantify Dfd levels using quantitative PCR assay, as not all discs are equally and strongly affected, making subtle changes in gene expression difficult to detect.

## Acknowledgments

We wish to thank Giorgia Giordani, Giuseppe De Candia, Laura De Grandis, Antonio Carusillo and Roberto Ciaccio for their help in preliminary phases of the project. Donatella Manzoni for excellent technical help and assistance. Manuela Voltattorni and Sabina Marianini for excellent help, advice and assistance with confocal microscopy. This study was supported by The Italian Association for Research on Cancer (IG15182 and IG24341 to G.P.).

## Author contributions

R.B., S.M., G.P., G.M. designed the experiments, S.M., R.B., S.S., E.D.G., M.S., S.K.Z., N.B., P.M. performed the experiments and S.M., R.B., A.G. and G.P. co-wrote the manuscript.

## Materials and methods

### Fly lines

Flies were reared on standard corn-yeast medium at 22°C. The following strains were used: *ey-Gal4* (Betz et al., 2001)*, UAS-dGFP* (gift from Gary Struhl) (Lieber et al., 2011), *UAS-dMyc* (BDSC #9674), *Max-RNAi* (BDSC #29328), *Dfd-RNAi* (BDSC #50792), *GFP-RNAi* (BDSC #41553 and BDSC #41555), *Myc^G0354^* (BDSC #11981), *Max^1^* (gift from Peter Gallant) (Steiger et al., 2008), *Max-rescue* (gift from Peter Gallant) (Steiger et al., 2008), *HDAC1-RNAi* (BDSC #33725), *HDAC1^04556^* (BDSC #11633), *Smr-RNAi* (BDSC #27068), *Smr^G0060^*(BDSC #11653). *UAS-MYCN* was generously donated by Daniela Grifoni (unpublished). Unless otherwise specified, both male and female animals were used in this study, to avoid gender bias.

### Phenotypic analysis of adult eyes

Animals were raised at the indicated temperatures (24 °C or 29 °C), and young adults (1-3 days old) were used for phenotypic analysis. Adult eyes were examined using a stereomicroscope, and eye defects were classified into high-dMyc, low-dMax and high-dMyc-low-dMax categories (see Results). For each genotype and temperature, > 100 flies were analyzed in several biological replicates. Phenotypic comparisons between experimental and control groups were performed using the Mann-Whitney U test.

### Fluorescent immunolabeling on eye imaginal discs

Wandering third instar larvae (wL3) were collected ∼120 hours after egg lay at 24 °C or ∼91 hours at 29 °C from eggs laid on yeast-supplemented agar medium. Larvae were dissected to expose the eye-antenna imaginal complex, fixed in 4% paraformaldehyde in 1× PBS for 20 min, and permeabilized in 0.3% Triton X-100 in 1× PBS (0.3% PBT) for 1 hour. Samples were blocked in 4% Normal Goat Serum (Jackson Immunoresearch) in 0.3% PBT for 20 minutes and incubated overnight at 4 °C with primary antibodies diluted in blocking solution. The following primary antibodies were used: monoclonal rat anti-Elav (1:100; 7E8A10 - DSHB), monoclonal mouse anti-Antp (1:400; 4C3 - DSHB), rabbit anti-Phospho Histone H3 (1:100, Upstate Biotechnology), mouse anti-Nubbin (1:50; 2D4 – DSHB). The next day, larvae were washed in 0.3% PBT, blocked for 20 min, and incubated with secondary antibodies for 2 hours. The following secondary antibodies were used: anti-mouse Cy3 (1:400, Jackson Immunoresearch), anti-rat FITC (1:400, Jackson Immunoresearch), anti-rabbit Cy5 (1:400, Jackson Immunoresearch). All the secondary antibodies were subtracted against the other hosts species to avoid cross-reactions. Cell nuclei were labelled with 2 μg/ml HOECHST (Sigma-Aldrich) in 1X PBS for 5 minutes. After three washes in 0.3% PBT, discs were isolated, mounted in Fluoromount-G (Electron Microscopy Sciences), dried overnight at room temperature, and imaged or stored at −20 °C.

### Imaging

Bright-field images of adult eyes and imago adults were captured as Z-stacks using a Nikon Eclipse T90i microscope at 10x magnification. Stacks were processed into single focused images using the Enhanced Deep Focus (EDF) module of NIS-Elements AR 3.10 software (Nikon).

Immunolabeled imaginal discs were imaged as Z-stacks with a Nikon Eclipse Ti2/A1R confocal microscope equipped with a PlanApo 40x objective and captured using NIS-Elements AR 3.10.

Maximum intensity projections generated in NIS-Elements AR 3.10 are shown in the figures, unless otherwise specified.

### Real-time quantitative PCR

Eye imaginal discs, separated from antennal discs, were dissected from 15 wL3 male larvae. Total RNA was extracted using TRI Reagent® (Sigma-Aldrich) according to the manufacturer’s instructions and resuspended in 15 µL RNase-free water. RNA was treated with the DNA-free™ Kit (Ambion, Thermo Fisher) and reverse-transcribed using iScript™ Reverse Transcription Supermix (Bio-Rad) with random primers. Reverse transcription was performed at 25 °C for 5 min, 42 °C for 30 min, and 85 °C for 5 min. Gene expression was quantified by real-time quantitative PCR using SsoAdvanced™ SYBR® Green Supermix (Bio-Rad) on a CFX96 real-time PCR system (Bio-Rad). The PCR program consisted of an initial denaturation at 95 °C for 30 s, followed by 30 cycles of 95 °C for 5 s and 60 °C for 30 s. A melting curve analysis was performed to verify amplification specificity. Relative expression of *dMyc* and *dMax* was calculated using the ΔΔCt method, with *RpL32n* and *Gapdh1* as reference genes. The primers used are listed below:

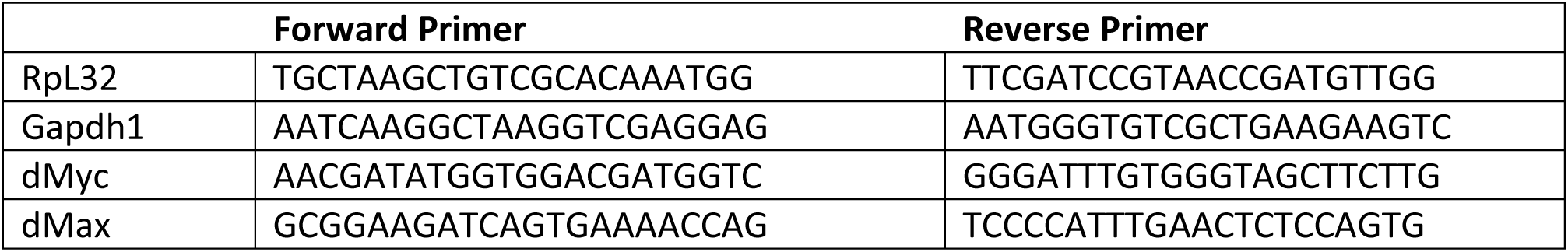

### Bulk RNA-sequencing assay

Larvae of the indicated genotypes (*2x ey-Gal4, 2x UAS-dMyc, 2x dMax-RNAi* or *2x ey-Gal4, 4x GFP-RNAi)* were generated from 2 hours egg lays on yeast-supplemented agar medium and reared at 24 °C until the wandering third instar (wL3) stage (∼120 h after egg lay). For each genotype, ∼60 eye imaginal discs were dissected, separated from the antennal discs, and collected in TRI Reagent® (Sigma-Aldrich) for RNA extraction. Three independent biological replicates of paired-end mRNA-sequencing libraries were prepared per genotype and sequenced.

Transcript abundance was quantified with Salmon v0.12.0 (Patro et al., 2017) against the *Drosophila melanogaster* reference transcriptome (dm6/BDGP6.32), using the --validateMappings and --gcBias options to improve mapping accuracy and correct for GC-content bias. Statistical analyses were performed in R 4.2.0 with Bioconductor 3.15. Transcripts’ quantifications were imported using the tximeta package (Love et al., 2020), and differential expression was assessed with DESeq2 (Love et al., 2020). Log2 fold changes were shrunk using the Bayesian estimator apeglm (Zhu et al., 2019) to improve accuracy with limited replicates. P-values were adjusted for multiple testing using the Benjamini–Hochberg false discovery rate (FDR), with adjusted p < 0.05 considered significant. Normalized counts were obtained through DESeq2 internal normalization followed by rlog transformation. Gene Ontology enrichment was performed with PANGEA ((Hu et al., 2023); https://www.flyrnai.org/tools/pangea/web/home/7227) on differentially expressed genes (DEGs), defined as those with | log2FC(high-dMyc-low-dMax/control) | > 1, adjusted p < 0.05. baseMean ≥ 14 (25th percentile cutoff) was used to exclude lowly expressed genes. The GO biological processes set was used.

