## Supplementary Figures for "MYC/MAX balance dictates cell progenitor fate by altering the HOX program in the *Drosophila* eye"

### Supplementary Figure 1

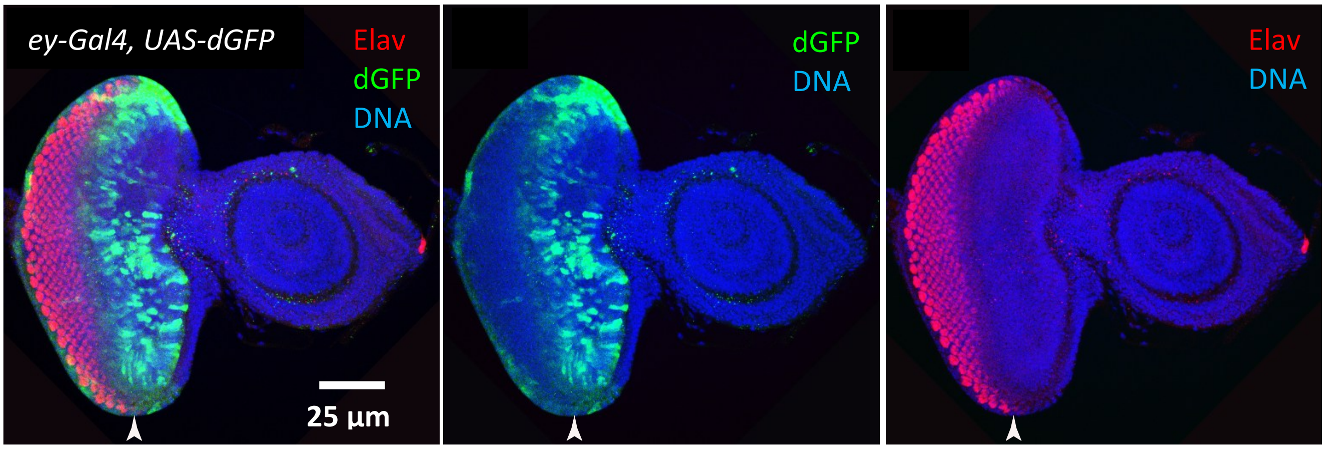


**Supplementary Figure 1** - **The *ey*-Gal4 driver is expressed by undifferentiated progenitors in the eye disc located anteriorly to the morphogenetic furrow.**

Immunolabeling assay on eye discs from wandering third instar (wL3) larvae expressing a destabilized GFP (*UAS-dGFP*) under the control of the *eyeless* promoter (*ey-Gal4*). GFP is in green, differentiating photoreceptors are labeled with anti-Elav (red), and nuclei are labelled with Hoechst (blue). Arrowheads mark the position of the morphogenetic furrow

### Supplementary Figure 2

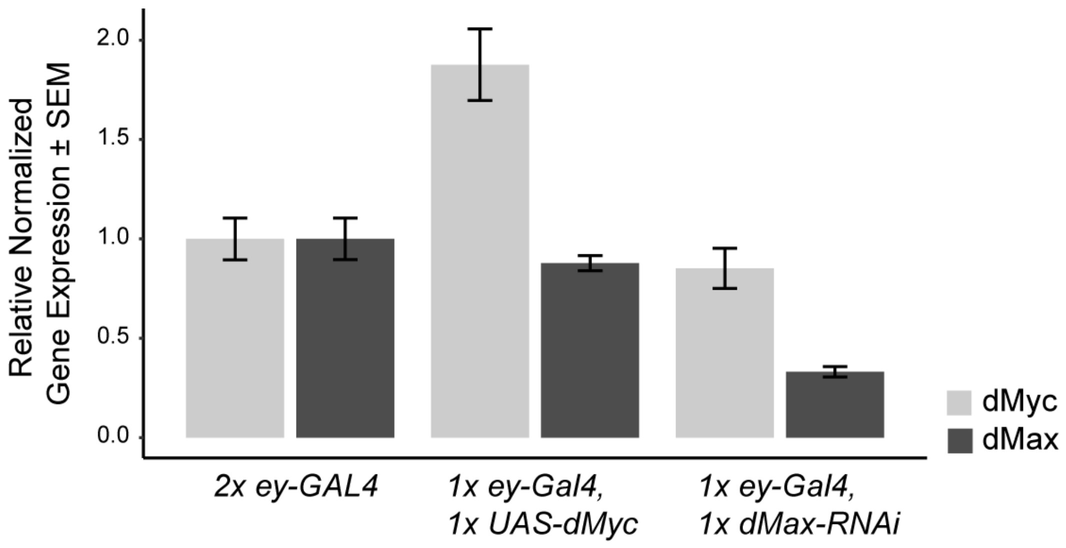


**Supplementary Figure 2 – Quantification of dMyc and dMax levels following dMyc overexpression and dMax downregulation**

qPCR assay on eye discs from third instar wandering L3 larvae (wL3, ~90 hour after egg lay at 29°C) comparing dMyc overexpression (*1x ey-Gal4, 1x UAS-dMyc*) or dMax downregulation (*1x ey-Gal4, 1x dMax-RNAi*) to controls (*2x ey-Gal4*). dMyc transcript levels approximately doubled in eye discs overexpressing dMyc, while dMax levels remained unchanged. Conversely, dMyc levels were unaltered in *dMax-RNAi* discs, whereas dMax transcripts were significantly reduced.

### Supplementary Figure 3

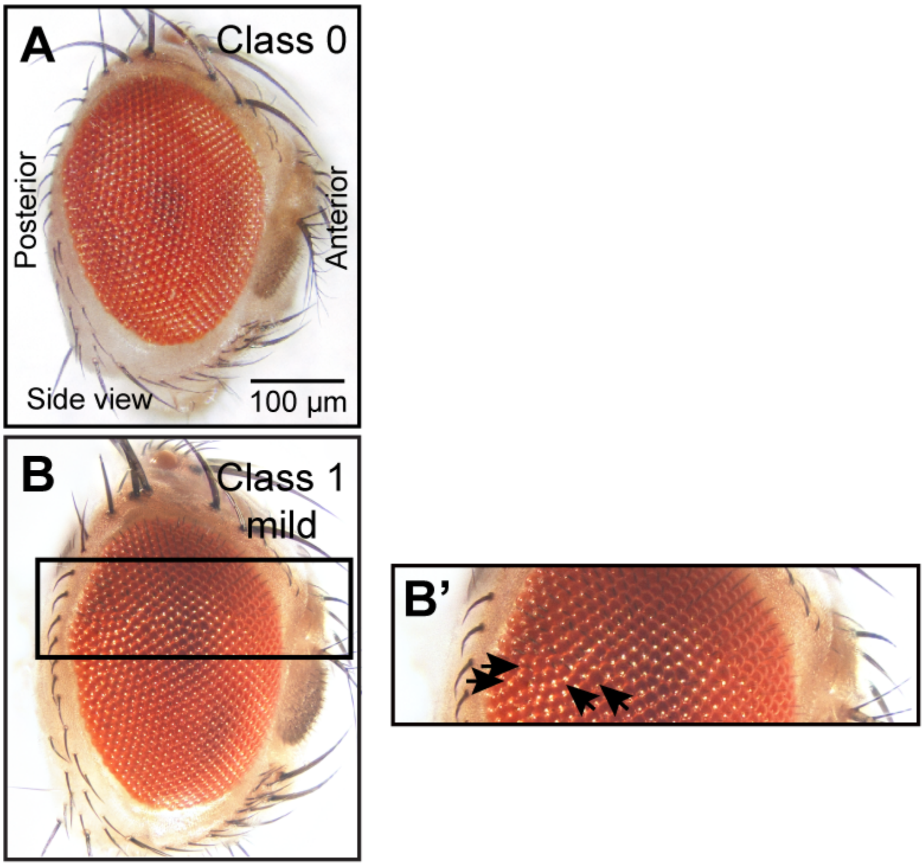


**Supplementary Figure 3 – Mild high-dMyc Class 1 phenotypic class in the adult eye.**

Adult eye bright-field images showing mild high-dMyc Class 1 phenotype observed upon overexpressing *Drosophila* Myc (*UAS-dMyc*) or the human MYCN (*UAS-MYCN*) in undifferentiated eye progenitors (*ey-Gal4*) **(B)** versus a Class 0, wild-type-like eye **(A).** Arrows point to eye defects: ommatidia misalignment mostly located in the posterior part of the eye. The box in **(B)** indicates the magnified view in **(B')**.

### Supplementary Figure 4

*2x ey-Gal4, 2x UAS-dMyc, 2x dMax-RNAi*

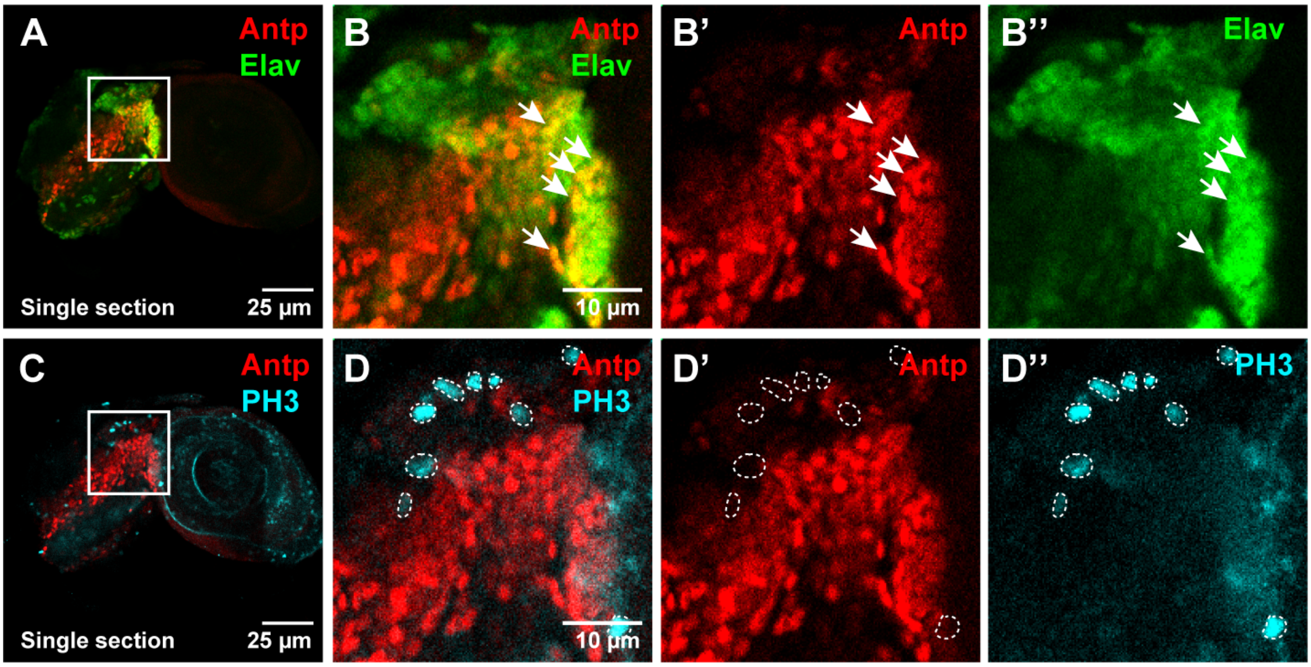


**Supplementary Figure 4 – High dMyc/dMax ratio induces ectopic expression of the wing HOX gene *Antennapedia* (*Antp*) in both Elav$^+$ and Elav$^-$ non-mitotic cells of the eye disc.**

Immunolabeling assay on eye discs from wandering third instar (wL3) larvae upon dMyc overexpression and dMax downregulation. **(A,C)** Single section confocal images of the eye disc shown in **Figure 4 B-B'''** with anti-Antp (red), anti-Elav (green) and anti-PH3 (cyan) labelling. **(B-B'')** High magnification of the region boxed in **(A)**, arrows indicate cells co-expressing Antp and Elav. **(D-D'')** High magnification of region boxed in **(C)** demonstrating absence of colocalization between Antp and PH3 signals.

### Supplementary Figure 5

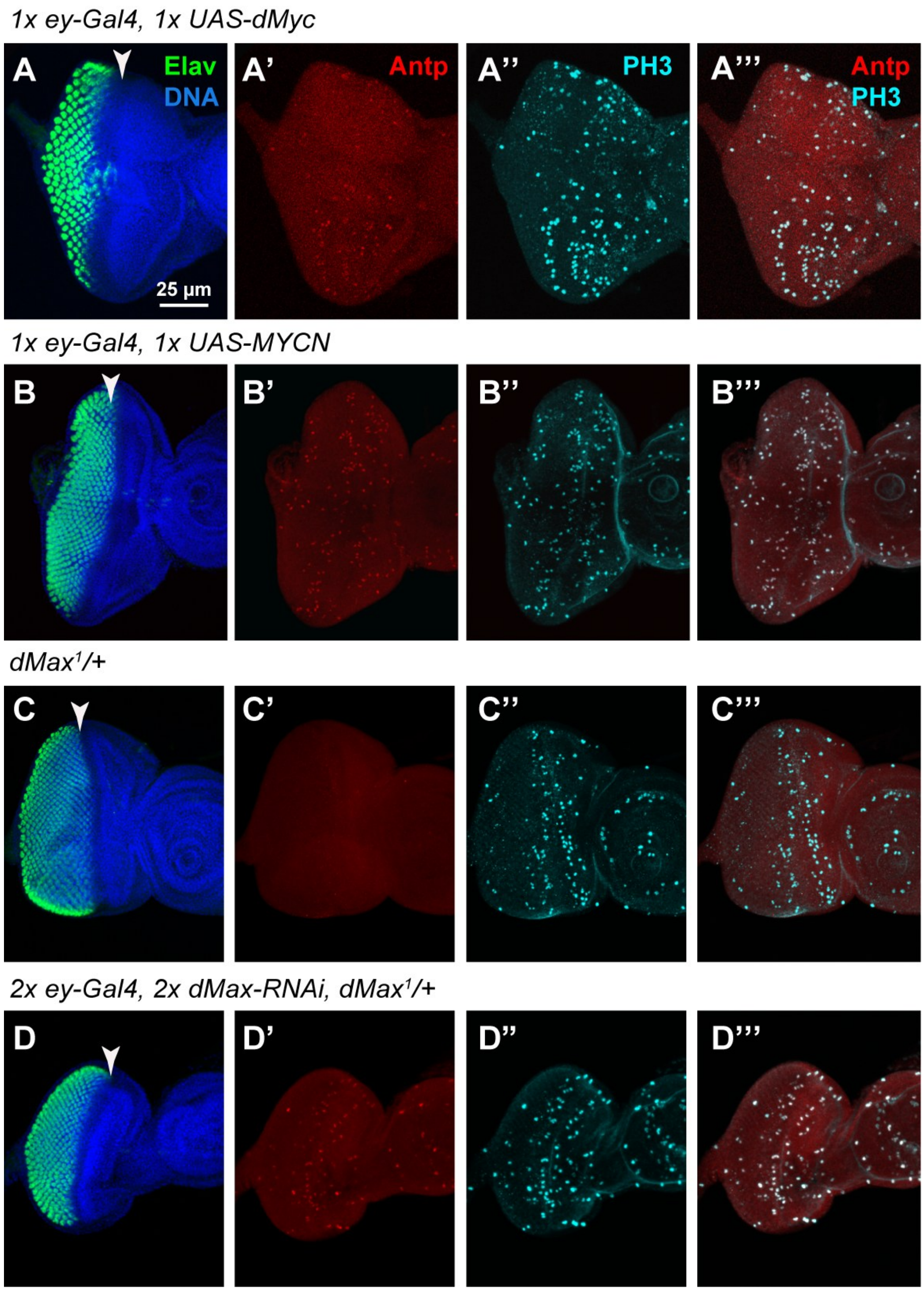


**Supplementary Figure 5 - The wing HOX gene *Antennapedia* (*Antp*) is ectopically expressed in eye discs upon high dMyc/dMax ratio.**

Immunolabeling assay on eye discs from wandering third instar (wL3) larvae upon **(A-A''')** dMyc overexpression, **(B-B''')** human MYCN overexpression or **(D-D''')** strong but viable dMax downregulation. Differentiating photoreceptors are labeled with anti-Elav (green), proliferating cells are labelled with anti-PH3 (Cyan), anti-Antp is in red, and nuclei are labelled with Hoechst (blue). Arrowheads mark the position of the morphogenetic furrow.

### Supplementary Figure 6

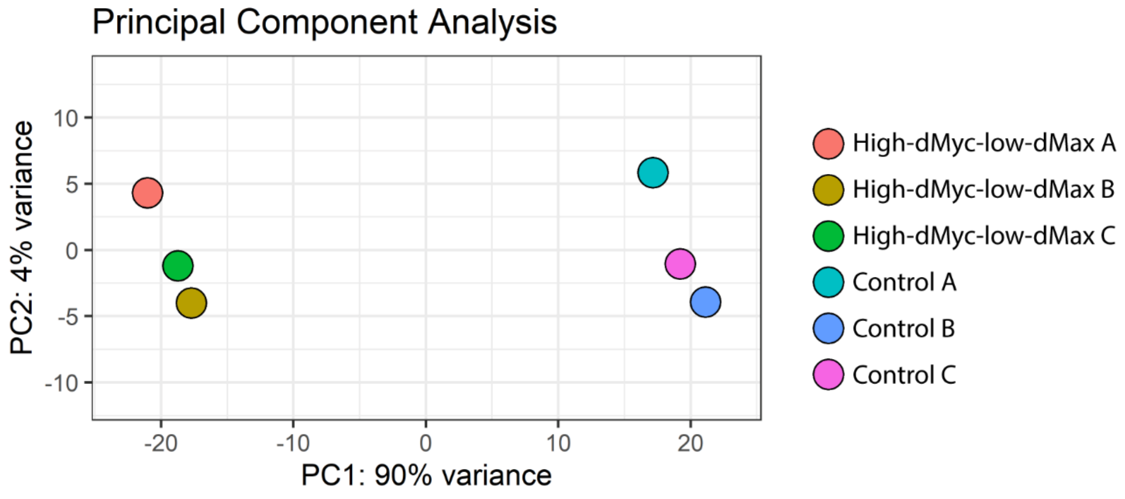


**Supplementary Figure 6** - **Principal Component Analysis (PCA) of bulk RNA-seq data.**

Principal component analysis performed on the normalized expression profiles of all the samples of the bulk RNA-seq assay: *2x ey-Gal4, 2x UAS-dMyc, 2x dMax-RNAi* (High-dMyc-low-dMax) and *2x ey-Gal4, 4x GFP-RNAi* (Control) wL3 eye imaginal discs, each done in triplicate (A, B and C). Each point represents an individual sample. Principal component 1 (PC1) and principal component 2 (PC2) are shown on the x- and y-axes, and explain 90% and 4% of the total variance in the dataset. The plot illustrates clustering of samples by genotype, indicating consistency across replicates of the same genotype

Supplementary Figure 7

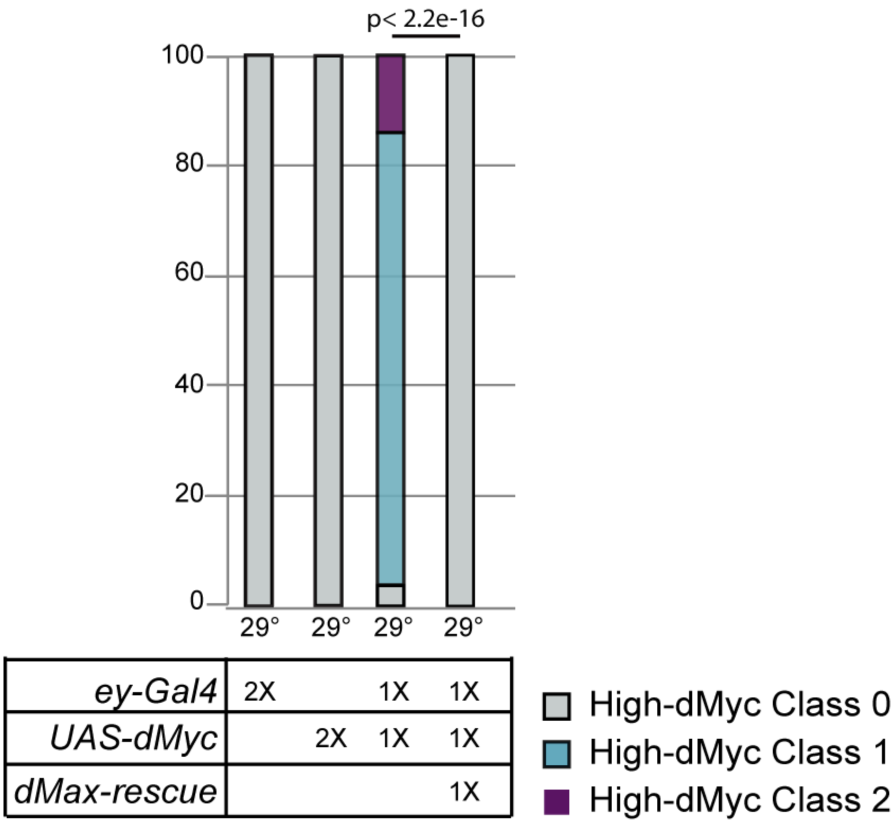


**Supplementary Figure 7 – Lowering dMax levels rescues the adult eye defects induced by dMyc overexpression.**

Quantification of the adult eye defects upon expressing the *UAS-dMyc* and *dMax-rescue* transgenes in the undifferentiated eye progenitors under the control of *ey-gal4*. Genotypes (1x = one transgene copy, 2x = two transgene copies) and rearing temperature are indicated below the bar chart. N ≥ 400. Statistics: Mann-Whitney.

### Supplementary Figure 8

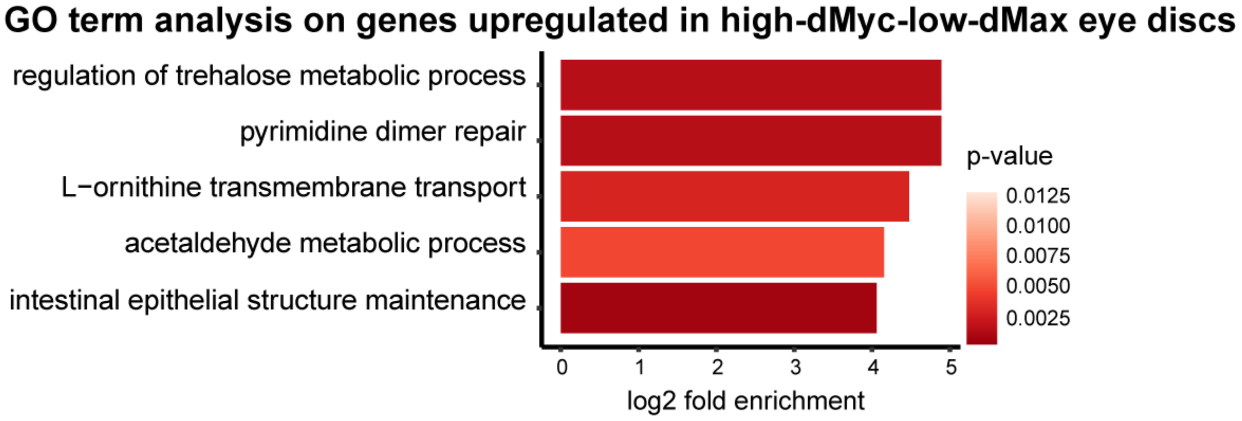


**Supplementary Figure 8 – High dMyc/dMax ratio induces a pro-oncogenic signature in the developing eye.**

Gene ontology (GO) analysis on the upregulated genes identified in the bulk RNA-seq assay on high-dMyc-low-dMax (*2x ey-Gal4, 2x UAS-dMyc, 2x dMax-RNAi*) and control (*2x ey-Gal4, 4x GFP-RNAi*) wL3 eye discs (~120 hour after egg lay at 24°C). Differentially expressed genes (DEGs) were defined based on |log2FC(high-dMyc-low-dMax/control)|>1, adjusted p-value < 0.05 and baseMean ≥ 14 (25th percentile cutoff). The top 5 GO terms are shown.

### Supplementary Figure 9

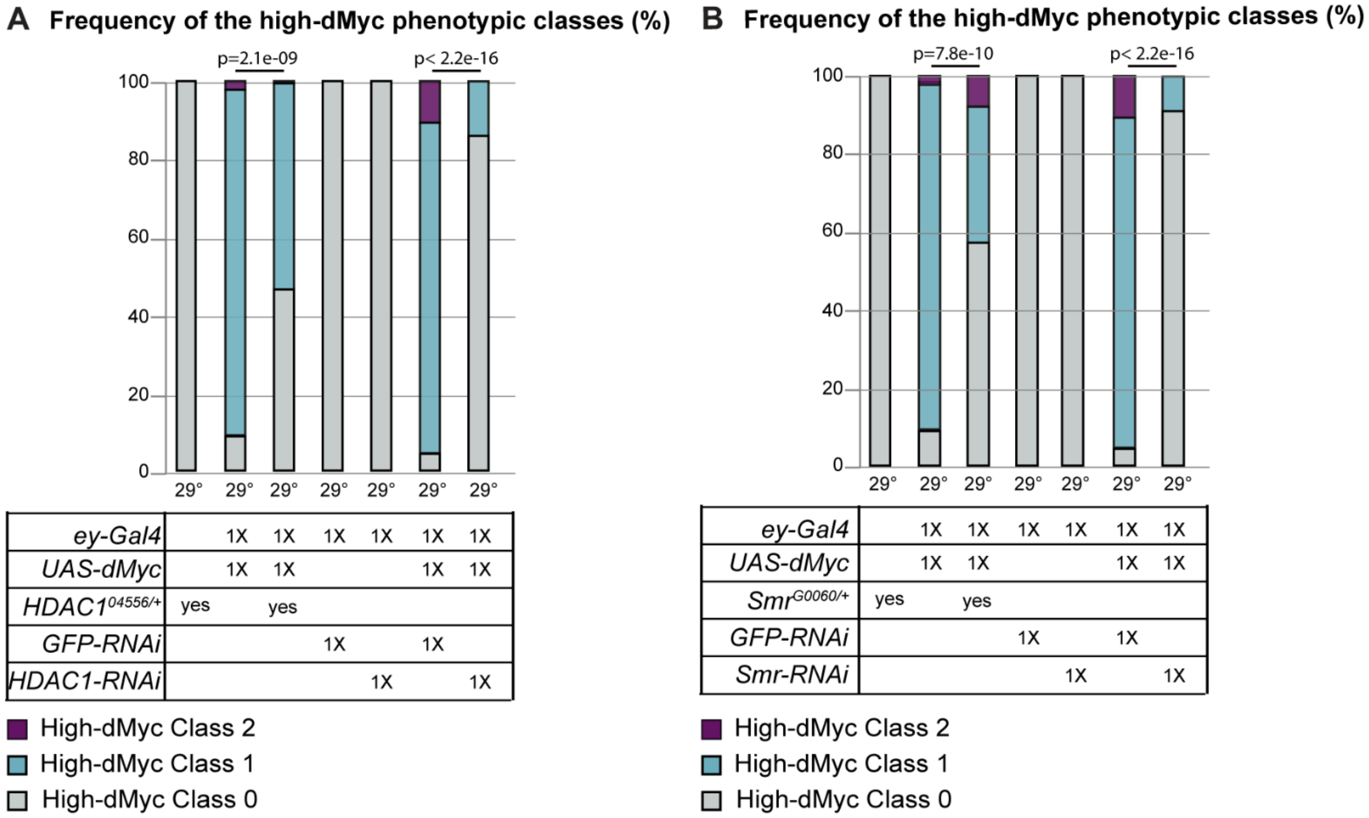


**Supplementary Figure 9 – Lowering the levels of the Histone deacetylase 1 (HDAC1) or Smrter (Smr) transcriptional co-repressors rescues adult eye defects induced by dMyc overexpression.**

Quantification of the adult eye defects upon expressing *UAS-dMyc* and silencing **(A)** HDAC1 or **(B)** Smr *via* RNAi in the undifferentiated eye progenitors under the control of *ey-gal4* or by using **(A)** HDAC1 or **(B)** Smr mutant alleles*.* Genotypes (1x = one transgene copy, 2x = two transgene copies) and rearing temperature are indicated below the bar chart. N ≥ 200. Statistics: Mann-Whitney.
